# Estimating the number of breeders from helminth larvae with genomic data

**DOI:** 10.1101/2023.08.25.554821

**Authors:** Tristan. P. W. Dennis, William Sands, Millicent Opoku, Alex Debrah, Linda Batsa, Kenneth Pfarr, Ute Klarmann-Schulz, Achim Hoerauf, Sabine Specht, Ivan Scandale, Lisa C. Ranford-Cartwright, Poppy H. L. Lamberton

## Abstract

Effective control of helminth infections requires the application of mathematical models to inform control efforts and policy, the development of product profiles for new drugs, and the monitoring of existing drugs for resistance. Key to the success of these approaches is accurately estimating the number of worms within a host, as well as distinguishing, in drug efficacy trials and monitoring, between adults surviving treatment and adults who have reinfected a host following drug treatment. In practice, observing adult worms is often extremely challenging in a patient, as many adult helminths are embedded deep in host tissues. Genetic approaches to infer kinship between larvae or eggs offer a solution to establish adult worm burdens, and to distinguish between treatment failure or treatment success followed by reinfection. Here, we use low-coverage whole-genome, and mitochondrial sequencing, of *Onchocerca volvulus* larvae to estimate the number of adults contributing to pools of offspring of known and unknown parentage. lcWGS reconstructs full-sibling relationships, resolving the number of unique adult worms contributing to a pool of offspring. Mitochondrial genotyping reconstructs maternal sib-ship, thus estimating the minimum number of adult females within a patient. Further development will improve these techniques for evaluating adult worm burden and trial outcome.

## Intro

Human helminthiases exert a devastating worldwide public-health burden. Many are neglected tropical diseases (NTDs), causing poor health, morbidity, and mortality, disproportionately for the world’s poorest people (1). Mass drug administration (MDA) with anthelmintic drugs remains the cornerstone of treatment for many NTDs. Monitoring and evaluation of these programmes is vital to support continued drug donation, to detect potential drug resistance, and to identify areas for improved interventions. The complex life cycle of many indirectly transmitted macroparasites makes elucidation of adult worm burden, the evaluation of control and drug trial success, and establishment of baselines of infection levels in pre-clinical trials difficult, as adult worms of many species (e.g. *Schistosoma mansoni* (2), *Wuchereria bancrofti* (3), *Onchocerca volvulus* (4), *Opisthorchis viverrini* (5) or *Trichuris trichiura* (6)) are either very difficult or impossible to retrieve from a patient. As such, inferring whether a trial macrofilaricide has successfully killed all adult worms, with any subsequent infection being due to reinfection, or whether treatment failure has been followed by repopulation of microfilariae (mf), or renewed reproduction and excretion of larvae from surviving adults, is impossible using current diagnostics. Moreover, estimating adult worm burden, a key parameter for assessing infection severity and individual contribution to community transmission, based on sampled offspring is problematic, as helminth larval abundance does not necessarily linearly predict adult worm burden, and instead often follows a density-dependent pattern (5,7,8), with adults only sporadically contributing to the larval pool at any given time (9). This gap introduces a considerable level of uncertainty for mathematical models – key tools for informing policy and control success.

*Onchocerca volvulus* is a filarial nematode parasite of humans, and is the agent of onchocerciasis, also known as river blindness, a disease characterised by blindness and debilitating, disfiguring skin disease. Onchocerciasis afflicts over 20 million people, with ∼1 million affected by blindness (10). The severe disease burden exerted by *O. volvulus* in Africa, South America and Yemen have led to concerted efforts to eliminate onchocerciasis. Since 1997, Community Directed Treatment with the microfilaricidal drug Ivermectin (Mectizan®) (CDTI) has been the mainstay of *O. volvulus* control and is the current World Health Organization (WHO) recommended strategy, administered as yearly MDA. Despite decades of CDTI, transmission and disease burden have remained stubbornly high in many regions (11–14). This difficulty necessitates both improved monitoring of existing CDTI, and the development and/or repurposing of anthelmintic drugs such as moxidectin (4), emodepside (15,16), oxfendazole (17) or Corallopyronin A (18). Evaluation of anthelmintics for *O. volvulus* is challenging. The adult worm is embedded in fibrous nodules, and whilst some of these nodules can be detected if near the skin’s surface, the adult worms themselves are accessible only through surgical nodulectomy - an invasive procedure. Diagnostics therefore rely on the counting of larval worms - microfilaria (mf) from skin biopsies, also an invasive procedure (albeit less so than a nodulectomy).

Genetic approaches for evaluating sib-ship and relatedness enable the estimation of the number of adults contributing to a sample of larvae (number of breeders, (*Nb)*), as well as establishing whether the adults contributing to serially sampled larvae have changed, e.g. if treatment has cleared the first set of parents, and a new set of parents have infected the patient. If full-siblings (FS) or maternal half-siblings (HS) between pre- and post-treatment samples are identified, treatment failure can be asserted due to the presence of surviving fecund females. Estimating the relationships and parentage of helminth larvae in the absence of adult information has been performed with multilocus genotyping methods such as microsatellite genotyping, with sib-ship and *Nb* reconstruction being performed most commonly with likelihood-based methods such as COLONY (19) for *Schistosoma mansoni* (20,21) and MASTERBAYES (22) for *Trichostrongylus tenuis*, methods which rely on well-validated microsatellite, SNP or multilocus sequence typing (MLST) panels.

Estimation of pairwise relatedness with genome data makes it possible to infer close familial relationships between individuals in the absence of validated multilocus data, and this is chiefly performed by estimating the proportion of alleles between two individuals that share 0, 1 or 2 copies identical-by-descent (IBD) (23). Whole-genome-sequencing is becoming increasingly affordable, especially with the advent of low-coverage WGS (lcWGS) approaches (24). lcWGS is particularly valuable if achieving high sequencing depth is problematic due to low-input and degraded samples - common for many helminth larvae (25). WGS is now a widely used tool for parasite molecular epidemiology (26,27), is possible from single helminth larvae (25), and can be used to determine relatedness (28,29). Another approach toward assigning parentage is mitochondrial genome sequencing. Mitochondria are inherited maternally, meaning that two individuals sharing identical, or near-identical mitochondrial genomes, can be assigned as probable maternal siblings. This is particularly useful in non-monogamous systems, such as many helminths including *O. volvulus (*which is thought to be polyandrous) (30), where a combination of relatedness estimation and mitochondrial genotyping can be used to identify maternal half-sibs. Due to higher copy number of the mitochondrial genome in the cell opposed to the nuclear genome, it is often easier to selectively amplify mitochondrial genomes than whole nuclear genomes for sequencing and analysis (24).

In this study, we perform a proof-of-concept with lcWGS, relatedness estimation, and mitochondrial genotyping in a complex indirectly transmitted macroparasite, *O. volvulus*. Our aims were to: 1) establish parentage and the *Nb* contributing to a sample of mf from which parentage is known, dissected from the uterus of gravid adult females, and 2) identify parentage of a sample of mf taken from a skin biopsy, of which parentage is unknown. Finally, we discuss the extent to which the approaches taken here may be more widely used toward estimating larval relatedness, adult worm burden, and treatment failure for any parasite where adults are inaccessible, but offspring, (eggs or larvae) can be obtained.

## Results

### 1. Parentage of intrauterine mf

We sequenced the genomes of 10 mf dissected from three adult females, from two separate human patients (**Figure 1**). The mean number of reads (‘X’) covering each position was 10.1 over the nuclear genome, and 47.6 over the mitochondrial genome. The exact number of per-sample reads is given in **Supplementary Table 1**. To calculate relatedness between each mf, we used genotype-likelihoods inferred from aligned reads as input for *NGSRelate*, which implements the framework for estimating relatedness and relationship described by (28). The inferred R0, R1, and KING kinship statistics from the intrauterine mf showed that mf collected from the same female worms/broods all formed a discrete cluster within the kinship coefficient thresholds for full- or half-sibling (28), (**Supplementary Figures 1A & 1B**). One pair from the same female fell outside of this (**Supplementary Figure 1A, 1B**). Based on these plots, we defined thresholds for full-sibling relationships at R0 ≤ 0.1, R1 ≥ 0.5, and KING ≥ 0.25, and thresholds for half-sibling relationships as R0 ≥ 0.1, R1 ≤ 0.5, and KING ≤ 0.25. These thresholds were then applied to our intrauterine mf data (**Supplementary Figure 2**).

**Figure 1:**
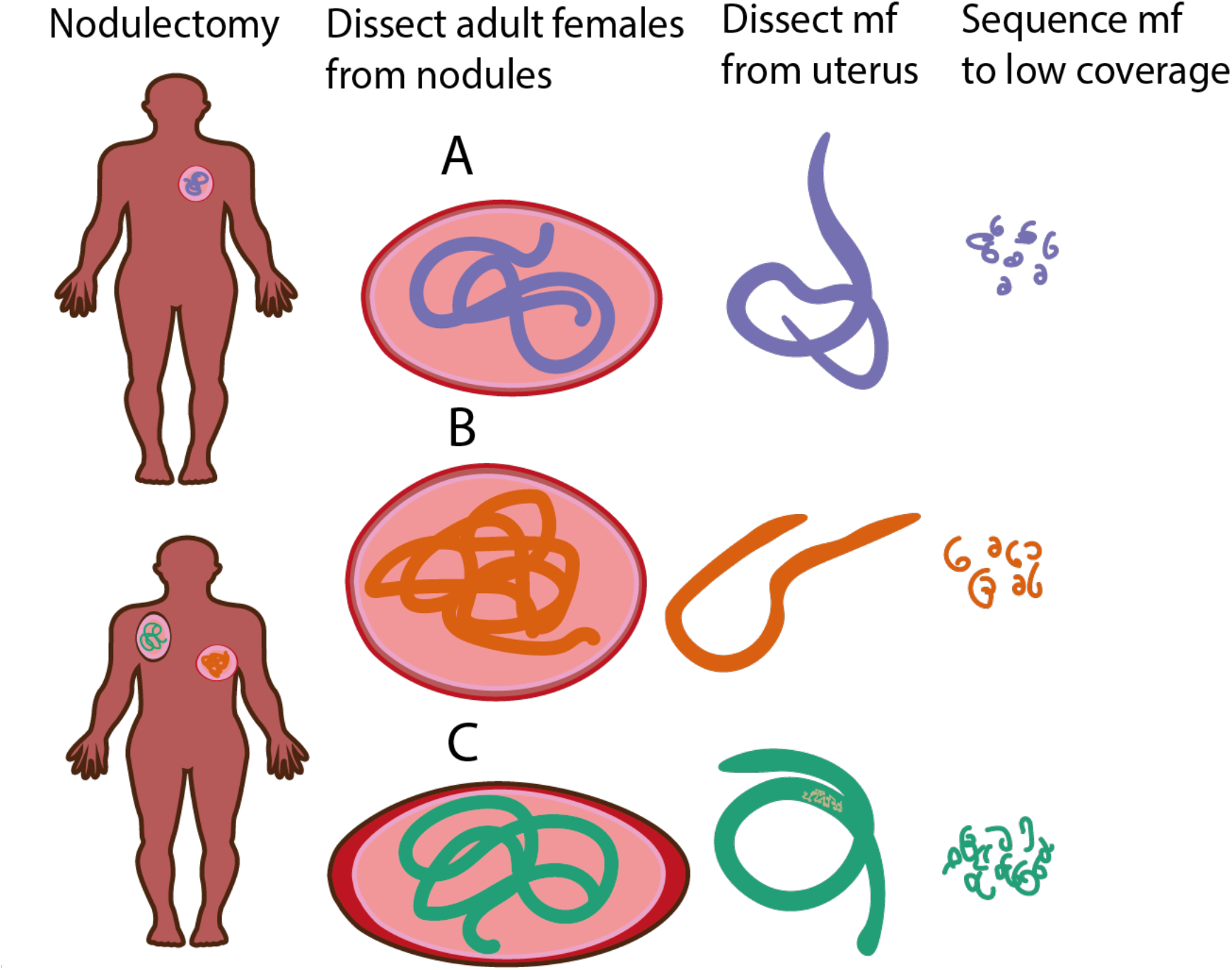
Schematic of *Onchocerca volvulus* microfilariae (mf) dissections from known parents. (Results Section 1) 10 mf were dissected from the uteri of 3 adult female worms, derived from 3 nodulectomies from 2 patients. mf were whole-genome-sequenced.

To further evaluate whether this method reconstructed sib-ship according to our expectations, we drew networks from the relatedness data and plotted them (**Figure 2**). We found that individuals dissected from the same female were at least full or half siblings. However, we found that the assignments also predicted half-sibling relationships to mf dissected from different females. As it is difficult to resolve relationships accurately below FS without nonsensical results (28,31), we discarded our HS estimations and retained only FS relationships (**Figure 2**). The network showed four FS groups, implying three female worms, and four male worms, contributed to the mf sample.

**Figure 2:**
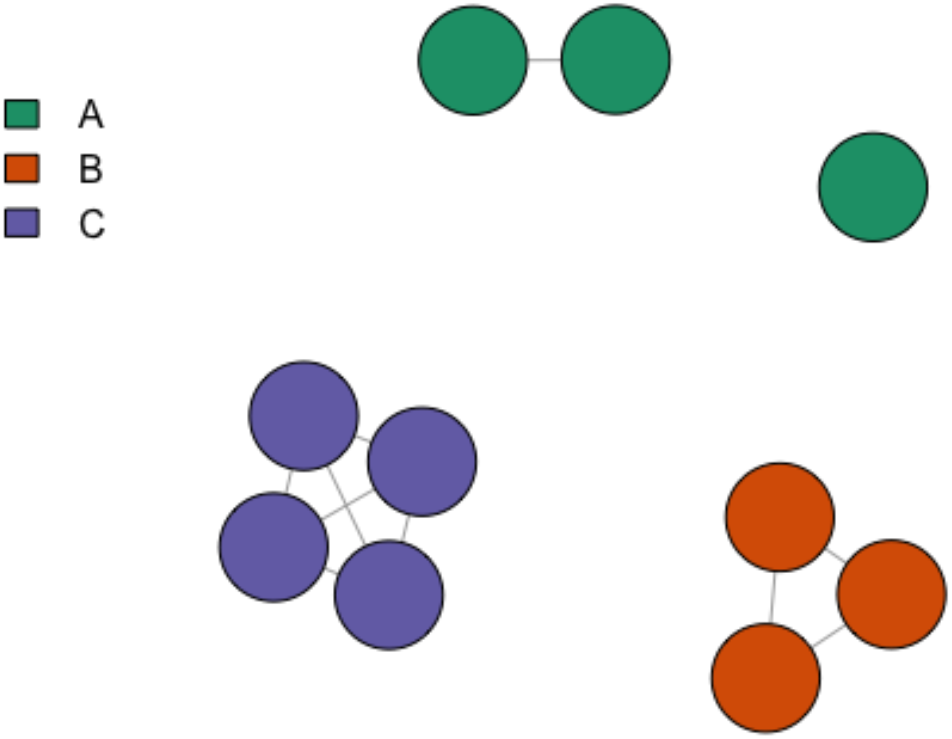
Network of full-sibling relationships between intrauterine *Onchocerca volvulus* microfilariae. Nodes indicate intrauterine *O. volvulus* microfilariae and edges indicate full-sibling relationships. Nodes are coloured by known maternal parentage.

We extracted mitochondrial genome *data in silico* from the intrauterine mf WGS data and used them to build a minimum-spanning network (**Figure 3**). The network showed a high-level of within-brood sequence similarity, with members of the same brood (having been dissected out of the same female uterus) being a maximum of three SNPs different to one another. This method reconstructed mitochondrial group membership with all offspring corresponding to the mother that they were dissected from (**Figure 3**). When the number of mitochondrial groups was considered alongside the number of full-sibling groups inferred from the intrauterine mf sample, we estimated that seven unique parents (three females and four males) had contributed to this sample of 10 mf.

### 2. Maternal assignment of skin-snip microfilariae from the same patient

To infer the number of breeders contributing to a sample of mf *for which the parentage was unknown*, we attempted to sequence the genomes of 39 mf from two skin snips taken from the left and right trochanter of a single patient. (**Figure 4**). Due to uneven and low nuclear genome coverage (**Supplementary Table 2**) (mean 4.1), we were unable to obtain reliable relatedness data for these samples. However, mitochondrial genome coverage across all samples was sufficient to genotype whole mitochondria genomes (mean depth-of-coverage 216.1X) (**Supplementary Table 2**). The medium-spanning network disclosed five major mitochondrial groups (**Figure 5**) indicating at least five adult females contributing to the 39 microfilariae for which we successfully obtained sufficient mitochondrial genome data from this patient. There was no structuring in mf haplotype by location (left or right-hand trochanter) **(Figure 5)**.

**Figure 4:**
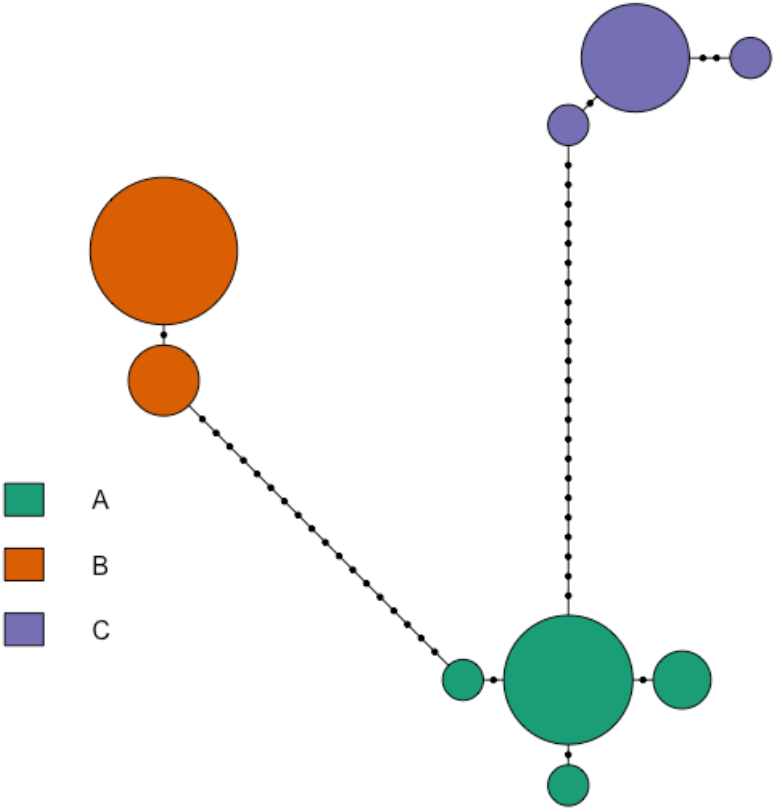
Medium-spanning network of intrauterine *Onchocerca volvulus* microfilariae mitochondrial haplotypes. Colour indicates shared known maternal parentage. Dots on connecting lines indicate the number of single nucleotide polymorphisms (SNPs) between haplotype groups.

**Figure 5:**
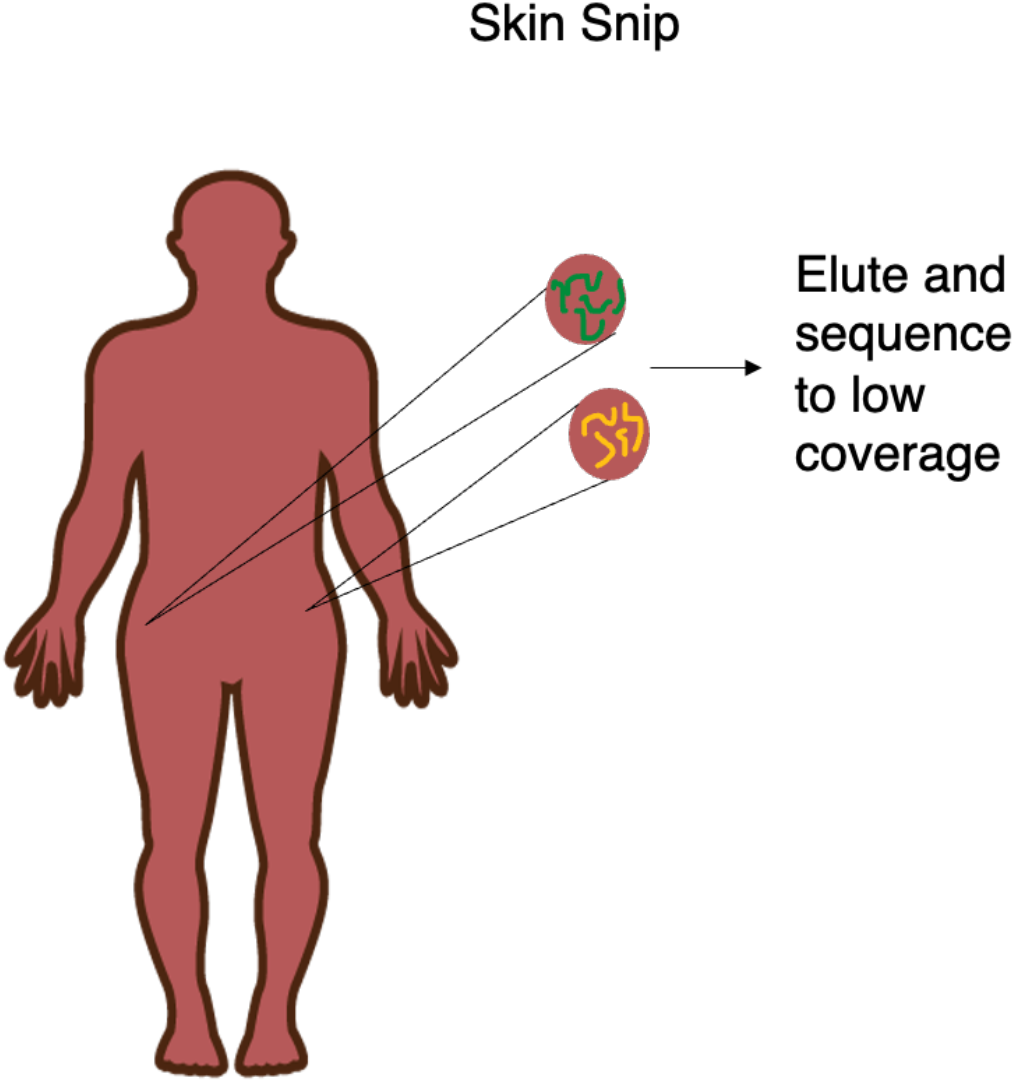
Schema of identifying parents of mf for which parentage is unknown. (Results Section 2) 39 mf were eluted from two skin-snips, from the left and right trochanter of a single patient.

**Figure 5:**
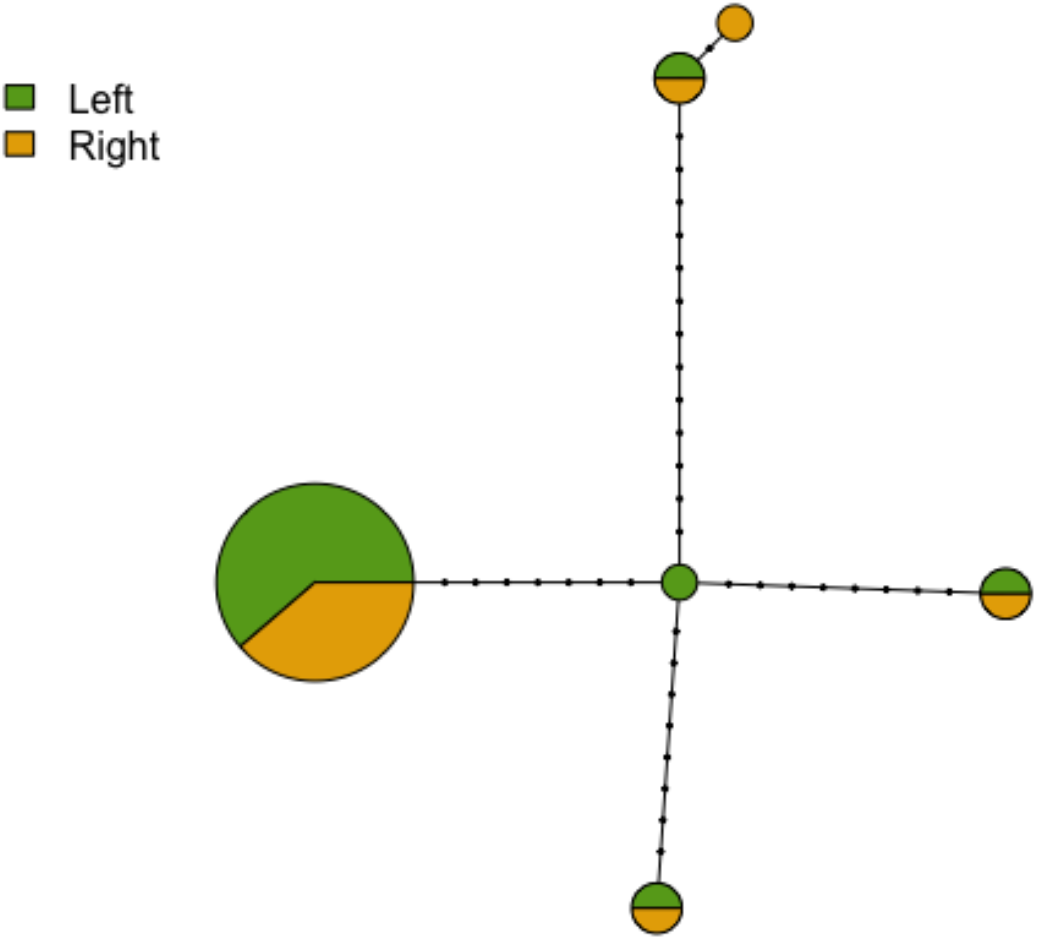
Medium-spanning network of mitochondrial haplotypes of skin-snip collected *Onchocerca volvulus* microfilariae from two locations on the body of a single patient. Pie colours indicate the proportion of samples taken from the left or right trochanter of the patient. Dots on connecting lines indicate the number of single nucleotide polymorphisms (SNPs) between mitochondrial haplotype groups.

## Discussion

For many helminth parasites, adult worms are not readily available for diagnosis, either at all from living patients (e.g. *Schistosoma spp*.), or only through invasive surgery (e.g. *O. volvulus)*, or inefficient adult worm excretion methods (e.g. *Ascaris, Opisthorchis*). Despite many transmission models and drug efficacy measurements being based on estimated adult numbers (32,33), diagnostics often only focus on the offspring. Here, we use intrauterine-collected mf from known *O. volvulus* adult females, as well as mf of unknown parentage collected from two skin-snips from one patient, to test methods to predict sib-ship relationships, maternal parentage and number of breeders (*Nb)*. We report the outcome of estimating helminth parentage using low-coverage WGS and relatedness estimation, as well as mitochondrial genotyping. For the intrauterine mf, we were successfully able to infer full-sibling relationships (including possible contribution of multiple fathers to the mf pool (30)), and, in the absence of successful half-sibling inference, we combined relatedness data with mitochondrial haplotypes to infer the *Nb* contributing to the offspring sample. For the skin-snip mf, we were hampered by degraded, low-input DNA leading to uneven genome coverage and unsuccessful relatedness estimation. However, good coverage over the mitochondrial genome enabled us to infer a minimum of five females contributing to the offspring samples.

Identification of shared parentage and *Nb* in parasitic helminths is a vital tool for discerning adult worm burdens, as well as for distinguishing between treatment failure followed by repopulation and treatment success followed by reinfection. This has the potential to inform the outcome of macrofilaricide clinical trials (e.g. (16,34)), as well as the establishment of key parasite infection dynamic parameters, such as the minimum adult worm burden and density-dependence, with the assistance of statistical models (e.g. (35,36)). Both relatedness estimation and mitochondrial genotyping have the potential to help address these questions in *O. volvulus* and other indirectly transmitted parasites. For *O. volvulus*, a chromosome level genome assembly has been generated (37), but no microsatellite, SNP or MLST panels exist for parentage analysis. Generating these resources typically require laboratory systems (20), as well as time, money, and access to samples. For pathogens for which laboratory life cycles are problematic to achieve, such as *O. volvulus*, and for which samples are in limited supply – for a variety of different reasons, including lack of funding, or difficult-to-obtain samples - generating panels of markers is difficult. Thus, methods such as ours, that rely on low-coverage WGS to infer relatedness, may be particularly suitable for studying infection dynamics and drug trial outcomes. Furthermore, once siblings have been excluded, unrelated samples can be used for parasite population genomic analysis without introducing biases from siblings. This method can easily be generalised to several parasite systems, if a reference genome assembly has been developed. Reference genomes are becoming increasingly common for even the most neglected of human helminthiases. However, methods for SNP discovery such as RAD-Seq also offer a potential stopgap for relatedness estimation in the absence of a reference genome, using relatedness methods such as SEQOUIA (38).

The sequencing effort reported here aimed for 5X coverage per-sample, which cost £30 per-sample (around $36 US at the time of writing). Options for improving the cost-effectiveness include performing downsampling experiments to establish the minimum depth-of-coverage required to identify kinships. Simulations suggest that *NGSRelate* may be robust to 1X coverage (39). Moreover, library preparation costs can be greatly reduced by adapting commercial library preparation kits (e.g. dilution) (24,40).

Another possibility toward reducing cost and increasing ease of access is to restrict parentage analysis in this form to the resequencing of mitochondrial genomes alone, which could be performed by designing long-range PCR primers to conserved mitochondrial regions (26). However, in the absence of population mitochondrial haplotype frequencies, it is impossible to determine whether two individuals with the same mitochondrial haplotype are truly maternal siblings, or whether their mothers are siblings, or if they have unrelated mothers with the same mitochondrial lineage. Here, our data indicate either there is homogeneous distribution of mf across an infected person despite adult worm location, or that there are multiple females with the same mitochondrial haplotype on either side of the body. Given extensive *O. volvulus* mitochondrial diversity and large population sizes (26), mitochondrial genotyping can at least establish the minimum number of females contributing to a pool of larvae, a valuable piece of information and a key parameter in models of helminth transmission (8,32,36), but it cannot determine the maximum number of females contributing to the offspring being assessed.

Despite technical improvements in single-larva DNA extraction and low-input library preparation (25), single-mf WGS is still technically challenging, with often unreliable results. The skin-snip mf, for example, sequenced very poorly, with highly heterogeneous coverage across the nuclear genome, perhaps indicative of degraded genomic DNA. Further work is needed to improve single-mf WGS sample storage and preparation both for the employment of this approach for use in epidemiological investigations and clinical trials, but also for improving *O. volvulus* population genomic analyses which hitherto has only been possible with adult worms (41,42). Population genomics based on mf would make evolutionary analysis of *O. volvulus* much easier and more ethical, as it removes the need for surgical nodulectomies to access adult worms, although the invasiveness of skin-snips remains a problem.

Ultimately, development of this method to the point where it can be reliably used in clinical trials will require optimisation of sample storage, DNA extraction and library preparation techniques, as well as more extensive testing against samples of mf for which parentage is known, so that the variation in the range of kinship coefficients corresponding to a given relationship and the error rates can be quantified. This is possible by performing further sequencing studies where adult worms are available.

## Conclusion

The decreasing cost of WGS, combined with technical and theoretical advances that make it possible to infer relationships between helminth larvae from low-coverage WGS data, are facilitating the development of techniques that enable interrogation of fine scale helminth infection dynamics, as well as identifying treatment failure, versus repopulation from surviving adult worms. This is important because many adult helminths, such as *O. volvulus* are, often inaccessible, with their offspring being comparatively easy to sample. Here, we employed lcWGS, relatedness estimation, and mitochondrial genotyping to identify parentage, maternal parentage and *Nb*, showing that, when sample data quality is sufficient, we are able to identify the number of males and females contributing to a sample of offspring and, even when nuclear genome data quality is poor, the minimum number of females may still be estimated. This method, when developed further has potential use in investigating the outcome of macrofilaricide trials for *O. volvulus*, as well as for further probing multiple parasite infection dynamics, in systems for which validated marker panels have not yet been developed.

## Materials and Methods

Data (sequencing metadata, relatedness data, mitochondrial group data) were parsed and analysed in *R* version *4*.*0*.*2*, (43), particularly using the *dplyr, gpplot2*, (44), *RColorbrewer* (45)and *cowplot* (46) packages.

### Sample Collection

#### Part 1

Onchocercomata containing adult female worms were surgically removed from patients as part of the field study “Comparison of doxycycline alone vs doxycycline plus rifampicin in their efficacy against onchocerciasis’ in Ghana in 2009.

#### Part 2

Skin-snips were collected in Oct 2019. All infected patients presented at least one palpable nodule. For Mf analysis, two skin biopsies of 1±3 mg were taken from the buttocks using a corneoscleral (Holth) punch. Each biopsy was immersed in 100 ml of 0.9% NaCl solution in a well of a microtiter plate. The skin biopsies were incubated overnight at room temperature to enable mf to emerge. The solution was then transferred onto a slide for microscopic examination. The biopsies were weighed using an electronic balance and mf load was calculated per mg skin. The skin biopsies and the solution containing the free mf were then stored at -20 degrees C for subsequent processing and analyses.

### Sample Processing, DNA Extraction and Sequencing Library Preparation

#### Part 1: Intrauterine mf

10 mf were dissected from three gravid *O. volvulus* adult females, obtained by surgical excision from two nodules. (**Figure 1**) Adult females were washed once in 70% ethanol and three times in PBS. Insulin needles (29G) were used to dissect mf from the adult female uterus. Individual mf were pipetted into a 96 well-plate, and DNA extracted using the Beckman Coulter Agencourt DNAdvance gDNA purification kit following the manufacturer’s instructions. DNA was eluted into 35μl. The presence of *O. volvulus* was determined using O-150 primers (47). 26 μl of DNA were fragmented and made into libraries using the New England Biolabs NEBNext Ultra FS II kit with unique dual barcodes. Libraries were pooled, and each pool size selected using a Sage Science Blue Pippin with a range of 150-500bp. Further QC was performed on an Agilent Bioanalyzer 2100 High-Sensitivity Chip.

#### Part 2: Skin-snip mf

39 mf were dissected into PBS with insulin needles from two skin-snips, one of the left and one of the right trochanter of a single patient. (**Figure 5**). The remaining processing followed the methods used for the intrauterine mf.

The intrauterine mf and skin-snip mf were then sequenced in two separate runs on an Illumina NextSeq Nano and an Illumina NovaSeq 6000, aiming for 13^e^6 paired-end reads (10X mean nuclear genome coverage of 2x75 paired-end reads) and 3.3^e^6 (5X coverage of 2x150 reads) respectively.

### Mapping and Genotype-Likelihood Calling

Demultiplexed reads were trimmed using *Trimmomatic* v0.39 (48), and aligned to the *O. volvulus* reference genome vWBPS15 (37) using bwa-mem (49). PCR duplicates were identified and removed with *GATK* (50). Samtools was used for SAM/BAM file manipulation, and calculation of mapping statistics (51). BAM files were used as input for ANGSD (52), which was used to calculate genotype likelihoods with the following filters: depth > 3, mapQ > 20, max-missingness <20% of individuals, Hardy Weinberg Equilibrium (HWE) p-value 1e-6. See **Supplementary Tables 1 and 2** for per-sample information, and [https://github.com/tristanpwdennis/onchogenome] for code.

### Relatedness Estimation

The final genotype likelihood file was used as input for the *NGSRelate* software (39), which infers, amongst others, the measures of relatedness: *R0, R1* and *KING-Robust-kinship*, as previously described (28,29,39). The relationship between pairs of full siblings was based on previously defined thresholds. All analyses downstream of *NGSRelate* were performed in R *4*.*0*.*2* (28,29) (see https://github.com/tristanpwdennis/onchogenome). R0/R1 and R1/KING-Robust-kinships were plotted with *ggplot2*. Graphs of related individuals were plotted in *iGraph* (53).

### Mitochondrial Genotyping

Reads mapping to the *O. volvulus* mitochondrial genome were extracted from the BAM alignments with samtools. SNPs were called using *Freebayes* (54) with a ploidy setting of 1, and filtered in *bcftools* (51), based on depth > 3 reads, maximum missingness < 0.25, QUAL > 30. The resulting filtered, multisample VCF file was converted into a FASTA alignment with bcftools *bcf-consensus*. The alignment was used as an input for a minimum-spanning network in PEGAS (55), in R. Clusters were defined based on the maximum level of divergence within-brood < 3 SNP differences.

## Ethics Statements

Onchocercomata containing adult female worms were collected as part of a field study “Comparison of doxycycline alone vs doxycycline plus rifampicin in their efficacy against onchocerciasis” in Ghana (ISRCTN68861628). Approval for the study was obtained from the Committee on Human Research, Publications and Ethics of the School of Medicine and Dentistry of the Kwame Nkrumah University of Science and Technology (Ref: CHRPE/KNUST/KATH/22-07-08), Kumasi, Ghana and the Ethics Committee of the University of Bonn, Medical Faculty (Nr. 186/08). Skin-snip collection was approved by the Committee on Human Research, Publications and Ethics of the School of Medicine and Dentistry of the Kwame Nkrumah University of Science and Technology (Ref: CHRPE/KNUST/KATH/22-07-08), Kumasi, Ghana, Research Ethics Committee of the Liverpool School of Tropical Medicine as well as the Ethical Committee of the University Hospital of Bonn, Germany.

## Data accessibility

All raw read data used in this study have been deposited in ENA under study accession PRJEB63468. All the code (analysis code as an RMarkdown file, sequence alignment/processing wrapping as shell wrappers), sequencing metadata, relatedness data, ANGSD output and mitochondrial genome alignments are available at https://github.com/tristanpwdennis/onchogenome.

## Acknowledgements

Many thanks to James Cotton and Thomas Crellen for helpful discussions and feedback on the manuscript.

## Supplementary Figures

**Supplementary Figure 1:**
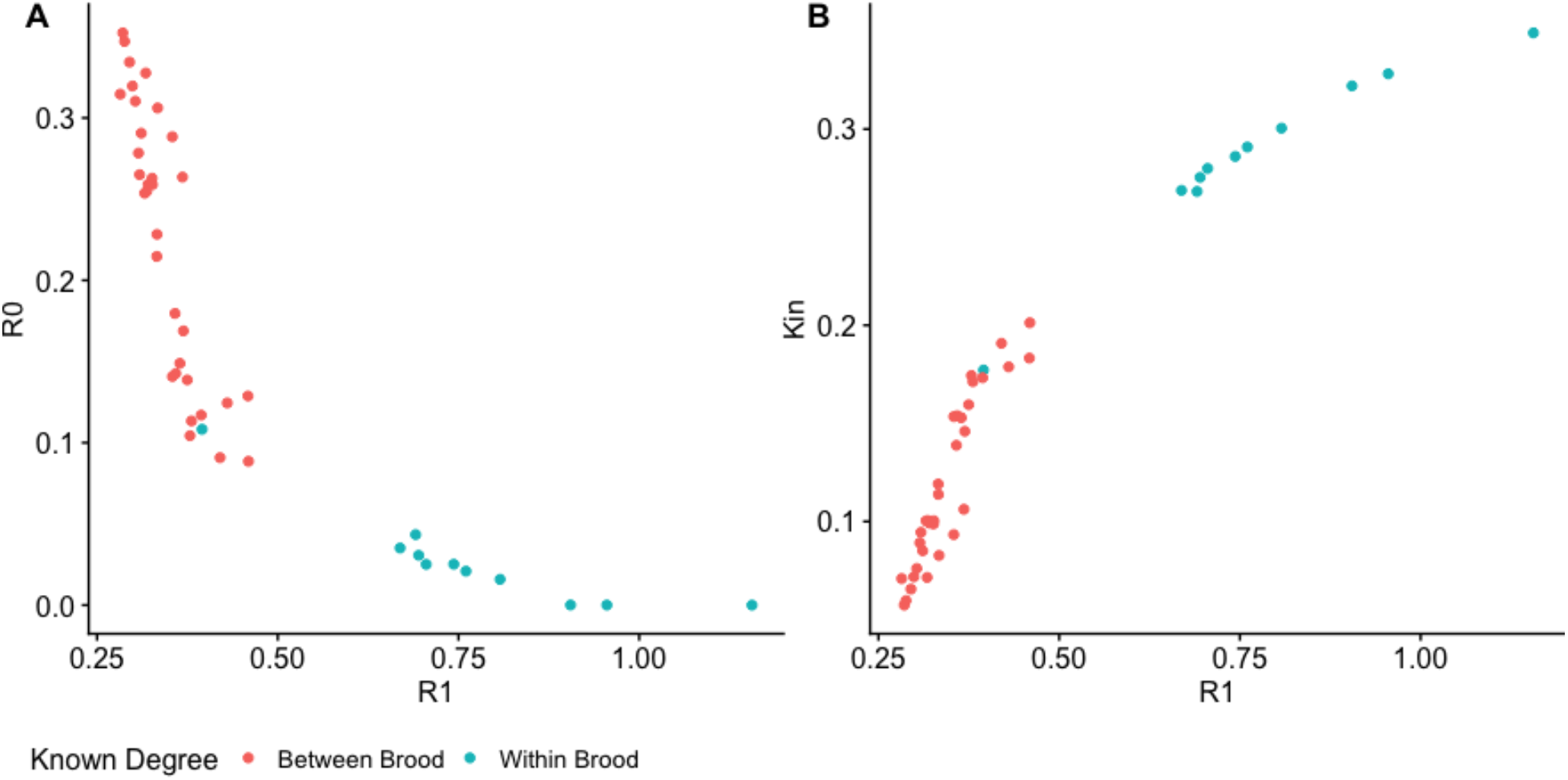
Inferred kinship statistics for R0, R1 and KING-robust-kinship for pairs of intrauterine *Onchocerca volvulus* microfilariae, with points coloured by whether individuals in a pair were sampled from the same (within brood) or different (between brood) *O. volvulus* adult female worms.

**Supplementary Figure 2:**
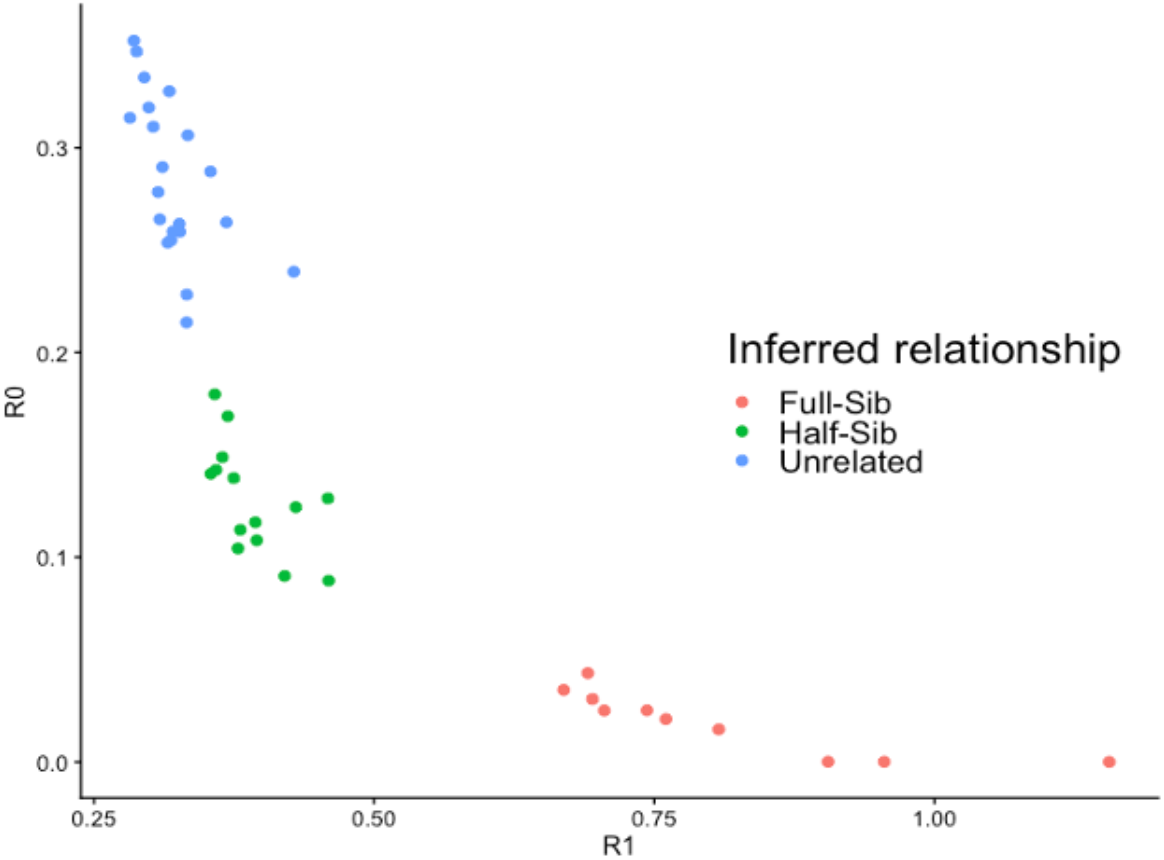
Kinship statistics R0 and R1 for pairs of intrauterine *Onchocerca volvulus* microfilariae, coloured by the inferred relationship using thresholds for full-sibling (Full-Sib) relationships at R0 < 0.1, R1 > 0.5, and KING > 0.25 and thresholds for half-sibling (Half-Sib) relationships as R0 > 0.1, R1 < 0.5, and KING < 0.25, informed from Figure 1.

